# Iota-carrageenan prevents the replication of SARS-CoV-2 on an in vitro respiratory epithelium model

**DOI:** 10.1101/2021.04.27.441512

**Authors:** Augusto Varese, Ana Ceballos, Carlos Palacios, Juan Manuel Figueroa, Andrea Vanesa Dugour

**Affiliations:** Instituto de Investigaciones Biomédicas en Retrovirus y SIDA (INBIRS), Universidad de Buenos Aires, Buenos Aires, Argentina; Respiratory Research Group, Instituto de Ciencia y Tecnología Dr. César Milstein - (Consejo Nacional de Investigaciones Científicas y Técnicas CONICET- Fundación Pablo Cassará), Saladillo 2468, Ciudad de Buenos Aires, C1440FFX, Argentina

**Keywords:** SARS-CoV-2_1_, COVID-19_2_, iota-carrageenan_3_, respiratory epithelium_4_, antivirals_5_

## Abstract

There are, except for remdesivir, no approved antivirals for the treatment or prevention of SARS-CoV-2 infections. Iota-carrageenan formulated into a nasal spray has already been proven safe and effective in viral respiratory infections. We explored this antiviral activity in Calu-3, a human respiratory model cell line. A formula of iota-carrageenan and sodium chloride, as a nasal spray, already approved for human use, effectively inhibited SARS-CoV-2 infection in vitro, providing a more substantial reference for further clinical studies or developments.

## 1 Introduction

The severe acute respiratory coronavirus 2 (SARS-CoV-2) is responsible for the currently ongoing pandemic coronavirus disease (COVID-19), counting more than 144.878.978 confirmed cases and more than 3.075.042 deaths worldwide by April 23, 2021 (Dong et al., 2020). There are still no adequate therapeutic or preventive medicines for COVID-19; repurposing established medications with recognized safety profiles is a possible approach for preventing or treating the disease and shortening the time-consuming drug development stages.

During the first days of the infection, the virus replicates mainly in the nasal cavity and the nasopharynx; therefore, nasal sprays with antiviral activity would reduce the viral load in these cavities.

Marine-derived polysaccharides, such as carrageenans, are a family of linear sulfated polysaccharides extracted from red seaweeds, widely used as thickening agents and stabilizers for food. Besides these properties, the iota-carrageenan demonstrated antiviral activity against several viruses, including respiratory viruses such as human rhinovirus, influenza A H1N1, and common cold coronavirus (Grassauer et al., 2008; Leibbrandt et al., 2010; Morokutti-Kurz et al., 2015). Iota-carrageenan inhibits virus infection mainly based on its interaction with the surface of viral particles, preventing them from entering cells and also trapping the viral particles released from the infected cells. It has also been shown that their inhibitory activity also relies on affecting the viral replication cycle at different steps, like entry and genome replication, and additionally activates the host’s antiviral immune response (Gomaa and Elshoubaky, 2016; Chen et al., 2020; Hans et al., 2020).

Iota-carrageenan formulated into a nasal spray has already been proven safe and effective in the common cold treatment (Koenighofer et al., 2014). Based on these observations, the hypothesis has been raised that a nasal spray with iota-carrageenan could be effective against SARS-CoV-2. It has recently been described that iota-carrageenan has activity against the SARS-CoV-2 virus and its Spike Pseudotyped Lentivirus (SSPL) in Vero E6 cell culture (Bansal et al., 2020; Morokutti-Kurz et al., 2020; Song et al., 2020). The Vero E6 cell line, originally derived from African green monkey kidney, is deficient for interferon-α (IFNα) and -β (IFNβ) due to genetic deletions, for instance highly susceptible to a vast number of different viruses, like measles virus, rubella virus, arboviruses, adenoviruses, influenza, and some coronavirus, including SARS-CoV-2 (Osada et al., 2014; Barrett et al., 2017; Banerjee et al., 2020).

Various studies have proposed the need to study SARS-CoV-2 infection in human respiratory epithelium, in order to get closer to the central target tissue of the disease in patients (Holwerda et al., 2020). Calu-3 is a non-small-cell lung cancer cell line that grows in adherent culture and displays epithelial morphology. This cell line is considered a sensitive and efficient preclinical model to study human respiratory processes and diseases (Zhu et al., 2010). Upon stimulation with viruses or environmental toxins, the Calu-3 cell line synthesizes and releases different cytokines, including IL-6 (Zhu et al., 2008), which play a central role in the inflammatory cascade associated with more severe COVID-19 (Gubernatorova et al., 2020).

The kinetics of SARS-CoV-2 infection show differences between Vero E6 and Calu-3 cells, most probably related to the differential expression of the angiotensin-converting enzyme 2 (ACE2) entry receptor and other facilitating molecules like the cellular serine protease TMPRSS2. Calu-3 cells express ACE2 on the apical membrane domain and are infected by SARS-CoV-2 via this route. The higher number of Vero E6 infected cells is also associated with the differences in viral entry pathway and the expression of pro-apoptotic proteins, which increased drastically in these cells but not in Calu-3 cells (Banerjee et al., 2020; Murgolo et al., 2021; Park et al., 2021; Saccon et al., 2021).

In this study, we assessed the effect of carrageenan as a viral infection inhibitor in an in vitro respiratory epithelium model, providing a set of data that support previous results and that it could be helpful against SARS-CoV-2 infection when applied in a formulation as a nasal spray.

## 2 Materials and Methods

### 2.1 Cells and Virus

African green monkey kidney Vero E6 cells (ATCC^®^ CRL-1586™) and human airway epithelial Calu-3 cells (ATCC^®^ HTB-55TM) were obtained from the American Type Culture Collection. The Calu-3 cells were cultured in Dulbecco’s modified Eagle’s medium (DMEM, Corning, NY, USA) containing 10% fetal bovine serum (FBS, Thermo Fisher Scientific, Waltham, MA, USA), 100 U/ml penicillin, and 100 μg/ml streptomycin (Thermo Fisher Scientific, Waltham, MA, USA). The Vero E6 cells were cultured in complete minimal essential medium (c-MEM) (Corning, NY, USA), supplemented with 5% FBS (Thermo Fisher Scientific, Waltham, MA, USA). The cells were incubated in 95% air and 5% CO_2_ at 37°C.

The SARS-CoV-2 isolate was kindly provided by Dr. Sandra Gallegos (National University of Córdoba, Argentina). Viral master seed stock was prepared using T175 flasks of Vero E6 cells. Each flask was harvested on day two post-infection, and the supernatant was centrifuged twice at 220 x g for 15 minutes to remove cellular debris. The titer of virus stock was determined by plaque assay on Vero E6 cells and expressed as plaque-forming units per ml (pfu/ml). The experiments using the virus were carried out in BSL3 facilities from the School of Medicine at the University of Buenos Aires.

### 2.2 Preparation of sample formulations

Solutions with iota-carrageenan and sodium chloride were prepared using a sterile nasal spray for therapeutic use. All the formulations and placebos were prepared at Laboratorio Pablo Cassará S.R.L. (Argentina) under aseptic conditions. The composition of active and placebo formulations is depicted in Table 1.

**Table 1.** Composition of candidate nasal formulations (samples containing iota-carrageenan)

| Component | Sample | Vehicle |
| --- | --- | --- |
| Iota-carrageenan | 1.7 mg/mL | - |
| Sodium Chloride | 9 mg/mL | 9 mg/mL |
| pH adjusted to | 6.00 – 7.00 | 6.00 – 7.00 |

To determine antiviral efficacy of formulations by titer reduction assay, sample formulations were used at a final iota-carrageenan concentration of 600 μg/ml; 60 μg/ml, 6 μg/ml, 0.6 μg/ml and 0,06 μg/ml. An equivalent concentration of placebos was used for titer reduction assay as controls.

### 2.3 Viability cellular assays

Calu-3 cells were seeded in 96-well tissue culture microplates at 3×10^4^ cells/well, and incubated overnight at 37°C under 5% CO_2_. Then, the cells were treated or not with iota-carrageenan 600, 60, 6, 0.6, and 0.06 μg/ml or vehicle in culture medium for 48 h at 37°C. After incubation, cells were washed and treated with MTS/PMS (CellTiter 96® Aqueous Non-Radioactive Cell Proliferation Assay, Promega, USA).

### 2.4 Infection assays

In three independent experiments, Calu-3 cells were seeded in 96-well tissue culture microplates at 3×10^4^ cells/well. After 48 h of incubation at 37°C, treated or not with iota-carrageenan 600, 60, 6, 0.6, and 0.06 μg/ml or vehicle and 2 h later infected with SARS-CoV-2 (multiplicity of infection (MOI) = 0.01 and 0.1) in serum-free DMEM (Thermo Fisher) for 1 h at 37°C. Then, cells were washed and placed in culture medium for 48 h. After that, supernatants were harvested and stored at −80°C.

### 2.5 Viral titration

Vero E6 cells were seeded into 96-well microplates and grown overnight at 37°C under 5% CO2. Tenfold dilutions of virus samples from Calu-3 cells were added to monolayers of 80% confluent Vero cells at 37°C for 1 h. After incubation, the inoculum was removed, and monolayers were overlaid with DMEM. The cells were incubated at 37°C for 72 h and fixed using 4% formaldehyde. Finally, cells were stained with 0.1% crystal violet in 20% ethanol and counted. Virus endpoint titer was determined using the Reed-Muench formula and expressed as 50% tissue culture infectious dose (TCID50) per ml.

### 2.6 Statistical analysis

In the cellular viability assays, One-way ANOVA followed by Dunnett’s multiple comparisons test was performed, and in the infection assays, the results were analyzed statistically by two-way

ANOVA, both using GraphPad Prism version 9.1.0, GraphPad Software, San Diego, California USA, https://www.graphpad.com.

## 3 Results

The antiviral effects of iota-carrageenan on SARS-CoV-2 were tested in a dose-dependent manner. In the first set of experiments, Vero E6 cells were pre-treated with different iota-carrageenan concentrations (600 μg/ml to 0.06 μg/ml), and cell viability was quantified (Figure 1A). No difference in cell viability was observed in iota-carrageenan treated cells compared to vehicle-treated control cells.

**Figure 1.**
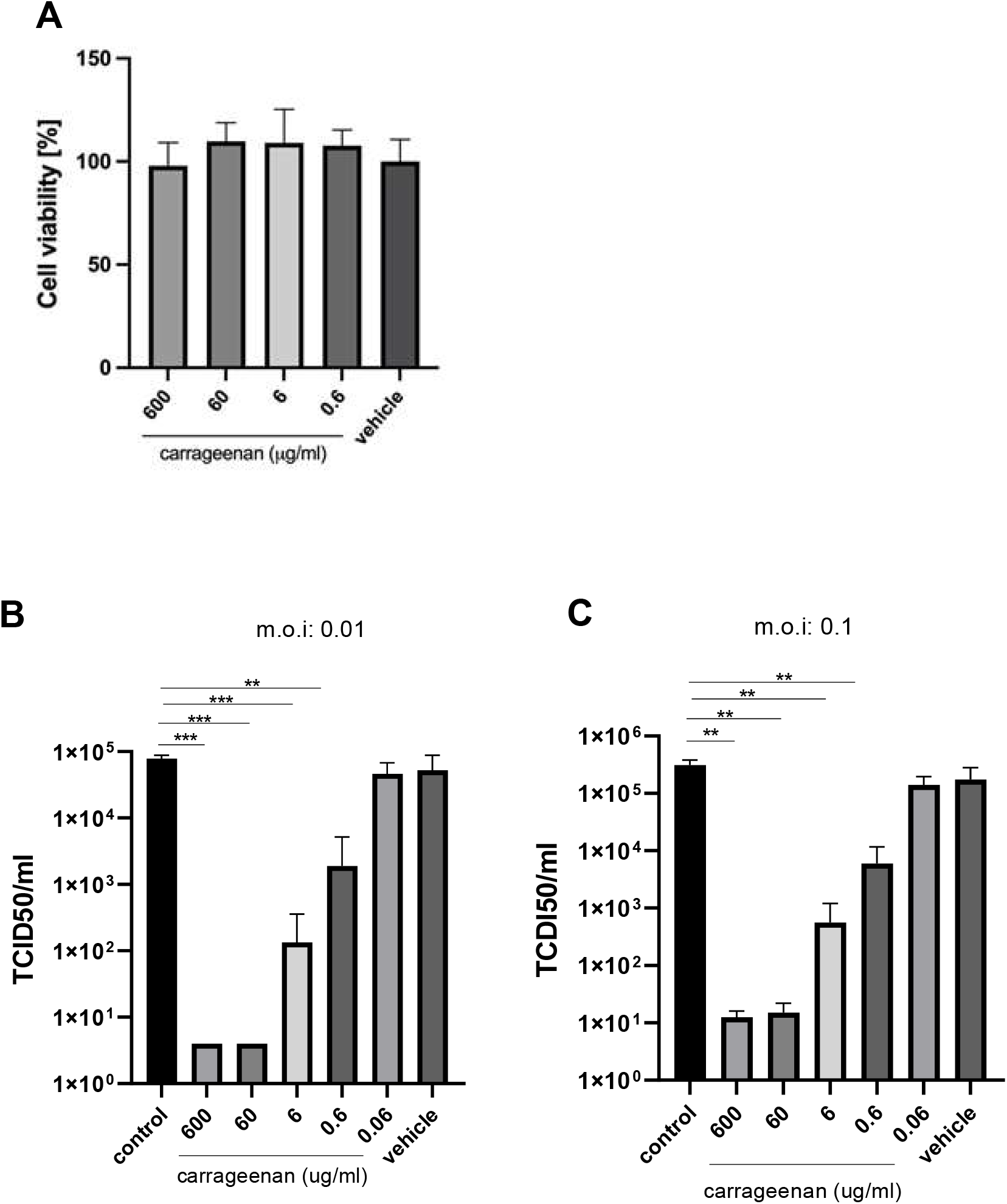
**A) Cellular viability assays.** Calu-3 cells were treated with iota-carrageenan or vehicle (600 μg/mL to 0 μg/mL) for 48 h at 37°C. After incubation, cellular viability was analyzed, and no statistically significant difference was found between the groups compared to the vehicle control group. Data are expressed as mean ± SD derived from three independent experiments. **B & C) Infection assays.** Calu-3 cells were pre-treated with iota-carrageenan or placebo (600 μg/mL to 0 μg/mL) for 1 h. After one hour of pretreatment, cells were infected, in two different conditions MOI: 0.01 (1B) and MOI: 0.1 (1C) with SARS-CoV-2 and incubated for 48 h in the presence of iota-carrageenan. Supernatants were harvested and virus yield. Data are expressed as mean ± SD derived from three independent experiments.

Next, Vero cells were pre-treated with iota-carrageenan (600 μg/ml to 0.06 μg/ml) for 2 h and then infected with SARS-CoV-2 (MOI: 0.01), after that, cells were washed to remove the viral inoculum, and fresh medium was added. Forty-eight hours later, supernatants were harvest. The SARS-CoV-2 production was evaluated by adding the supernatants to Vero E6 cells for 1 hour. After incubation, the inoculum was removed, and monolayers were incubated at 37 °C for 72 hours. Then cells were fixed and stained with crystal violet. Virus endpoint titer was determined by Reed-Muench formula and expressed as TCID50/ml. Our results showed that iota-carrageenan markedly inhibits SARS-CoV-2 production in a dose-dependent manner (Figure 1, B and C). No antiviral activity was only observed at the lowest concentration of iota-carrageenan (0.06 μg/ml). Lastly, there was no reduction in virus production with vehicle formulation, suggesting that the iota-carrageenan, not the sample excipient components, inhibited the SARS-CoV-2 replication (Figure 1B). Finally, Figure 1C shows that inhibition of viral production is also observed when pretreatment with carrageenan is applied to the cells, followed by an infection at a higher MOI (0.1).

## 4 Discussion

Calu-3 cell culture infection has been used as a model to evaluate the activity against SARS-CoV-2 of different drugs: suramin (Salgado-Benvindo et al., 2020), nafamostat (Yamamoto et al., 2020), exogenous interferon (Felgenhauer et al., 2020), chloroquine, and hydroxychloroquine (Hoffmann et al., 2020).

Recent research has shown that the Vero and Calu-3 cells results do not always coincide. One remarkable example is hydroxychloroquine’s antiviral activity when tested in Vero cells, which could not be reproduced on infected Calu-3 cells. These preliminary results, obtained in Vero cells, prompted the premature use of this drug to prevent or treat COVID-19 in patients. Still, the results of clinical trials carried out later have not shown efficacy, in accordance with what was observed in Calu-3 studies.

When considering Vero E6 as a model, it should consider their deficient expression ACE2 and TMPRSS2 and the non-specific endocytic viral uptake mechanisms responsible for viral entry in this cell line. Calu-3 cells, besides of the exposed before, are very similar to primary cultures of bronchial epithelium obtained by biopsy or surgery; it develops the characteristic tight junctions present in the respiratory epithelium, expression of cystic fibrosis transmembrane conductance regulator (CFTR) chloride channels, capacity for the secretion of mucus and proteins towards the apical end and exchange of water and electrolytes (Duszyk, 2001; Zhu et al., 2008).

In our research, we used dilutions of a commercially available iota-carrageenan spray. The concentrations found to be active in vitro, as shown in previous (Bansal et al., 2020), and in the current work, are those that would be achieved using the spray according to the approved dosage. These results are supporting the efficacy of this spray in the prevention of COVID-19 in a clinical trial recently carried out with frontline healthcare personnel exposed to SARS-CoV-2 (Figueroa et al., 2021).

## 5 Conclusion

In summary, our results confirm that a formulation of iota-carrageenan and sodium chloride available as a nasal spray effectively inhibited SARS-CoV-2 infection in vitro in human respiratory epithelial cell line culture, strengthening the hypothesis that a nasal spray with iota-carrageenan may be helpful in the prevention or treatment of COVID-19 and reinforces the interest in the development of clinical trials on this topic.

## 7 Author Contributions

AC, AVD, and JMF conceptualized the study. AC, AV, CP and AVD developed the methodology. JMF wrote and prepared the original draft. AC, AVD, and CP wrote, reviewed, and edited the manuscript. AC and AVD supervised the study. All authors contributed to the article and approved the submitted version.

## References

Banerjee, A., Nasir, J. A., Budylowski, P., Yip, L., Aftanas, P., Christie, N., et al. (2020). Isolation, Sequence, Infectivity, and Replication Kinetics of Severe Acute Respiratory Syndrome Coronavirus 2. Emerg Infect Dis 26, 2054–2063. doi:10.3201/eid2609.201495.

Bansal, S., Jonsson, C. B., Taylor, S. L., Figueroa, J. M., Dugour, A. V., Palacios, C., et al. (2020). Iota-carrageenan and Xylitol inhibit SARS-CoV-2 in cell culture. Biorxiv, 2020.08.19.225854. doi:10.1101/2020.08.19.225854.

Barrett, P. N., Terpening, S. J., Snow, D., Cobb, R. R., and Kistner, O. (2017). Vero cell technology for rapid development of inactivated whole virus vaccines for emerging viral diseases. Expert Rev Vaccines 16, 883–894. doi:10.1080/14760584.2017.1357471.

Chen, X., Han, W., Wang, G., and Zhao, X. (2020). Application prospect of polysaccharides in the development of anti-novel coronavirus drugs and vaccines. Int J Biol Macromol 164, 331–343. doi:10.1016/j.ijbiomac.2020.07.106.

Dong, E., Du, H., and Gardner, L. (2020). An interactive web-based dashboard to track COVID-19 in real time. Lancet Infect Dis 20, 533–534. doi:10.1016/s1473-3099(20)30120-1.

Duszyk, M. (2001). CFTR and lysozyme secretion in human airway epithelial cells. Pflügers Archiv 443, S45–S49. doi:10.1007/s004240100643.

Felgenhauer, U., Schoen, A., Gad, H. H., Hartmann, R., Schaubmar, A. R., Failing, K., et al. (2020). Inhibition of SARS-CoV-2 by type I and type III interferons. J Biol Chem, jbc.AC120.013788. doi:10.1074/jbc.ac120.013788.

Figueroa, J. M., Lombardo, M., Dogliotti, A., Flynn, L. P., Giugliano, R. P., Simonelli, G., et al. (2021). Efficacy of a nasal spray containing Iota-Carrageenan in the prophylaxis of COVID-19 in hospital personnel dedicated to patients care with COVID-19 disease A pragmatic multicenter, randomized, double-blind, placebo-controlled trial (CARR-COV-02). doi:10.1101/2021.04.13.21255409.

Gomaa, H. H. A., and Elshoubaky, G. A. (2016). Antiviral Activity of Sulfated Polysaccharides Carrageenan from Some Marine Seaweeds. International Journal of Current Pharmaceutical Review and Research 1, 32–34.

Grassauer, A., Weinmuellner, R., Meier, C., Pretsch, A., Prieschl-Grassauer, E., and Unger, H. (2008). Iota-Carrageenan is a potent inhibitor of rhinovirus infection. Virol J 5, 107. doi:10.1186/1743-422x-5-107.

Gubernatorova, E. O., Gorshkova, E. A., Polinova, A. I., and Drutskaya, M. D. (2020). IL-6: relevance for immunopathology of SARS-CoV-2. Cytokine Growth F R 53, 13–24. doi:10.1016/j.cytogfr.2020.05.009.

Hans, N., Malik, A., and Naik, S. (2020). Antiviral activity of sulfated polysaccharides from marine algae and its application in combating COVID-19: Mini review. Bioresour Technology Reports 13, 100623. doi:10.1016/j.biteb.2020.100623.

Hoffmann, M., Mösbauer, K., Hofmann-Winkler, H., Kaul, A., Kleine-Weber, H., Krüger, N., et al. (2020). Chloroquine does not inhibit infection of human lung cells with SARS-CoV-2. Nature 585, 588–590. doi:10.1038/s41586-020-2575-3.

Holwerda, M., V’kovski, P., Wider, M., Thiel, V., and Dijkman, R. (2020). Identification of an Antiviral Compound from the Pandemic Response Box that Efficiently Inhibits SARS-CoV-2 Infection In Vitro. Microorg 8, 1872. doi:10.3390/microorganisms8121872.

Koenighofer, M., Lion, T., Bodenteich, A., Prieschl-Grassauer, E., Grassauer, A., Unger, H., et al. (2014). Carrageenan nasal spray in virus confirmed common cold: individual patient data analysis of two randomized controlled trials. Multidiscip Resp Med 9, 57. doi:10.1186/2049-6958-9-57.

Leibbrandt, A., Meier, C., König-Schuster, M., Weinmüllner, R., Kalthoff, D., Pflugfelder, B., et al. (2010). Iota-Carrageenan Is a Potent Inhibitor of Influenza A Virus Infection. Plos One 5, e14320. doi:10.1371/journal.pone.0014320.

Morokutti-Kurz, M., Graf, P., Grassauer, A., and Prieschl-Grassauer, E. (2020). SARS-CoV-2 in-vitro neutralization assay reveals inhibition of virus entry by iota-carrageenan. Biorxiv, 2020.07.28.224733. doi:10.1101/2020.07.28.224733.

Morokutti-Kurz, M., König-Schuster, M., Koller, C., Graf, C., Graf, P., Kirchoff, N., et al. (2015). The Intranasal Application of Zanamivir and Carrageenan Is Synergistically Active against Influenza A Virus in the Murine Model. Plos One 10, e0128794. doi:10.1371/journal.pone.0128794.

Murgolo, N., Therien, A. G., Howell, B., Klein, D., Koeplinger, K., Lieberman, L. A., et al. (2021). SARS-CoV-2 tropism, entry, replication, and propagation: Considerations for drug discovery and development. Plos Pathog 17, e1009225. doi:10.1371/journal.ppat.1009225.

Osada, N., Kohara, A., Yamaji, T., Hirayama, N., Kasai, F., Sekizuka, T., et al. (2014). The Genome Landscape of the African Green Monkey Kidney-Derived Vero Cell Line. Dna Res 21, 673–683. doi:10.1093/dnares/dsu029.

Park, B. K., Kim, D., Park, S., Maharjan, S., Kim, J., Choi, J.-K., et al. (2021). Differential Signaling and Virus Production in Calu-3 Cells and Vero Cells upon SARS-CoV-2 Infection. Biomol Ther. doi:10.4062/biomolther.2020.226.

Saccon, E., Chen, X., Mikaeloff, F., Rodriguez, J. E., Szekely, L., Vinhas, B. S., et al. (2021). Tropism of SARS-CoV-2 in commonly used laboratory cell lines and their proteomic landscape during infection. Biorxiv, 2020.08.28.271684. doi:10.1101/2020.08.28.271684.

Salgado-Benvindo, C., Thaler, M., Tas, A., Ogando, N. S., Bredenbeek, P. J., Ninaber, D. K., et al. (2020). Suramin Inhibits SARS-CoV-2 Infection in Cell Culture by Interfering with Early Steps of the Replication Cycle. Antimicrob Agents Ch 64. doi:10.1128/aac.00900-20.

Song, S., Peng, H., Wang, Q., Liu, Z., Dong, X., Wen, C., et al. (2020). Inhibitory activities of marine sulfated polysaccharides against SARS-CoV-2. Food Funct 11, 7415–7420. doi:10.1039/d0fo02017f.

Yamamoto, M., Kiso, M., Sakai-Tagawa, Y., Iwatsuki-Horimoto, K., Imai, M., Takeda, M., et al. (2020). The Anticoagulant Nafamostat Potently Inhibits SARS-CoV-2 S Protein-Mediated Fusion in a Cell Fusion Assay System and Viral Infection In Vitro in a Cell-Type-Dependent Manner. Viruses 12, 629. doi:10.3390/v12060629.

Zhu, Y., Chidekel, A., and Shaffer, T. H. (2010). Cultured Human Airway Epithelial Cells (Calu-3): A Model of Human Respiratory Function, Structure, and Inflammatory Responses. Critical Care Res Pract 2010, 394578. doi:10.1155/2010/394578.

Zhu, Y., Miller, T. L., Singhaus, C. J., Shaffer, T. H., and Chidekel, A. (2008). Effects of oxygen concentration and exposure time on cultured human airway epithelial cells. Pediatr Crit Care Me 9, 224–229. doi:10.1097/pcc.0b013e318166fbb5.

